# A pan-cancer analysis of the frequency of DNA alterations across cell cycle activity levels

**DOI:** 10.1101/2020.02.24.962571

**Authors:** Arian Lundberg, Linda S. Lindström, Joel S. Parker, Elinor Löverli, Charles M. Perou, Jonas Bergh, Nicholas P. Tobin

## Abstract

**Background:** Pan-cancer genomic analyses based on the magnitude of pathway activity are currently lacking. Focusing on the cell cycle, we examined the DNA mutations and chromosome arm-level aneuploidy within tumours with low, intermediate and high cell cycle activity.

**Patients and methods:** Matching mRNA, gene mutational status, chromosomal arm-level aberrations and clinico-pathological data was assembled from pan-cancer studies of 9,515 patients with 32 different cancers. Cell cycle activity was estimated from mRNA data using the cell cycle score (CCS) signature. Barplots were used to visualise mutation and chromosomal aberration frequency within CCS subgroups. Kaplan-Meier and multivariable Cox-regression analyses were used to determine survival differences between CCS subgroups.

**Results:** Cell cycle activity varied broadly across and within all cancers. *TP53* and *PIK3CA* mutations were common in all CCS subgroups but with increasing frequency as cell cycle activity levels increased (*P* < 0.001). Mutations in *BRAF* and gains in 16p were less frequent CCS high tumours (*P* < 0.001). In Kaplan-Meier analysis, patients whose tumours were CCS Low had a longer PFI relative to intermediate or high (*P* < 0.001) and this significance remained in multivariable analysis (CCS intermediate: HR = 1.37; 95% CI 1.17 – 1.60, CCS high: 1.54; 1.29 – 1.84, CCS Low = Ref).

**Conclusions:** Cell cycle activity varies across and within cancers and whilst similar DNA alterations can be found at all activity levels, some notable exceptions exist. These data also demonstrate that independent prognostic information can be derived on a pan-cancer level from a simple measure of cell cycle activity.

## Introduction

The Nobel prize winning research of Hartwell [1], Nurse [2,3] and Hunt [4] in the nineteen seventies and eighties fundamentally changed our understanding of the cell cycle and provided broad insight into the molecules governing its regulation. These seminal discoveries have shaped our modern view of the cell cycle and its separation into four distinct phases commonly referred to as G1, S, G2 and M. Transitions between these phases are governed by the cyclin family of proteins along with their binding partners the cyclin dependent kinases (CDKs) [5]. Disruptions to the function of cyclin-CDK holoenzymes or other cell cycle pathway members can lead to impaired control over the cycle and sustained proliferation - a hallmark of cancer [6].

Large scale pan-cancer studies have sought to understand human malignancies at a molecular level through the integration of multiple high-throughput data types. This approach has yielded a number clinically relevant findings including the coalescence of lung squamous, head and neck, and some bladder cancers into a single pan-cancer subtype and the ability to classify tumours into prognostic subgroups at a pan-cancer level [7]. More recently, data from over eleven-thousand patients has shown actionable mutations in up to fifty-seven percent of tumours [8], a positive correlation between aneuploidy and cell cycle genes [9], and frequent co-alterations in the p53 and cell cycle pathways [10]. To date, the analysis of genomic aberrations in these studies have typically focused on all pan-cancer tumours at once [8], within subgroups of tumours that have clustered together on the basis of DNA, RNA and protein expression – termed the iClusters [8], or within tumours with a common genetic alteration such as chromosome 3p loss [9]. Given the varying degrees of oncogenic pathway activation/suppression across cancer types [10], we hypothesized that basing genomic analyses on the magnitude of pathway activity may also provide important biological information and clinical insight. In view of the fundamental biological role of the cell cycle in cancer and the frequent genomic alterations of its pathway members, it represents a compelling choice for a pathway activity-based analysis.

Here, in order to test our hypothesis, we compare the most prevalent genomic alterations in tumours with low, intermediate and high levels of cell cycle activity by integrating data from multiple genomic platforms in over nine-thousand tumours from The Cancer Genome Atlas (TCGA). Specifically, we examine gene expression levels, gene mutational frequency and chromosome arm-level alterations across pan-cancer tumours grouped into tertiles of cell cycle activity on the basis of our cell cycle score (CCS) gene signature [11,12]. Finally, we also determine the clinical relevance of this signature across and within cancer types using survival analyses including Kaplan-Meier graphs and multivariable Cox proportional hazards modeling adjusting for patient and tumour characteristics.

## Patients and Methods

### Study population and specimens

The Pan-Cancer Atlas (PanCanAtlas) project compared and contrasted genomic and cellular differences between tumour types profiled as part of TCGA. The project consists of 11,069 patients with primary tumours from 32 different cancer types, including Adrenocortical carcinoma (ACC), Bladder Urothelial Carcinoma (BLCA), Brain lower grade Glioma (LGG), Cervical squamous cell carcinoma and endocervical adenocarcinoma (CESC), Cholangiocarcinoma (CHOL), Colon adenocarcinoma (COAD), Esophageal carcinoma (ESCA), Glioblatoma multiforme (GBM), Head and Neck squamous cell carcinoma (HNSC), Kidney Chromophobe (KICH), Kidney renal clear cell carcinoma (KIRC), Kidney renal papillary cell carcinoma (KIRP), Liver hepatocellular carcinoma (LICH), Lung adenocarcinoma (LUAD), Lung squamous cell carcinoma (LUSC), Lymphoid Neoplasm Diffuse Large B-cell Lymphoma (DLBC), Ovarian serous cystadenocarcinoma (OV), Pancreatic adenocarcinoma (PAAD), Pheochromocytoma and Paraganglioma (PCPG), Prostate adenocarcinoma (PRAD), Rectum adenocarcinoma (READ), Sarcoma (SARC), Skin Cutaneous Melanoma (SKCM), Stomach adenocarcinoma (STAD), Testicular Germ Cell tumours (TGCT), Thymoma (THYM), Thyroid carcinoma (THCA), Uterine Carcinosarcoma (UCS), Uterine Corpus Endometrial Carcinoma (UCEC) and Uveal Melanoma (UVM).

From the original 11,069 patients, 9,515 were included in our study and reasons for exclusion were missing or no matching gene expression data (n = 795), copy number data (n = 498) or clinico-pathological information (n = 261). A CONSORT diagram showing the exclusion criteria for this study is shown in Supplemental figure 1. All clinical, gene expression, mutation and chromosome arm-level data from the PanCanAtlas study were taken from the publicly available database of the National Institutes of Health (NIH) (https://gdc.cancer.gov/about-data/publications/pancanatlas).

### mRNA data, clustering and the Cell Cycle Score (CCS)

Fully processed, batch corrected, RNA-sequencing data were accessed from NIH genomic data commons (GDC) database (https://gdc.cancer.gov). All data quality control, normalisation and gene level counts were performed by the PanCanAtlas investigators as described in the their original publication [13]. Integrative Cluster (iCluster) were also retrieved from the same publication. Cluster of cluster assignments (COCA) were performed by the pan-can investigators as described in Hoadley *et al.* [7], resulting in 32 different tumour clusters. Clusters with less than 20 tumours were excluded from further analysis. Cell Cycle Score (CCS) signature was applied as previously published [11,12]. Briefly, we extracted gene expression data from 433 of 463 signature CCS genes from all pan-cancer tumours and summed their values on an individual tumour basis to derive a single score of cell cycle activity for each sample. This continuous variable was further divided into tertiles in order to classify tumours as having Low, Intermediate or High levels of cell cycle activity on a broad, pan-cancer level. Cancer types where the pan-cancer CCS demonstrated independent prognostic information in multivariable Cox proportional hazard models were also assessed using within (intra-) cancer CCS tertiles: KIRC, LGG, LUAD, PAAD, SARC, UCEC and UVM.

### Mutational analysis

Fully processed mutational data derived from exome sequencing was taken from GDC database in a mutation annotation format file (MAF) (https://gdc.cancer.gov). All data quality control, processing and mutation calling was performed by the PanCanAtlas investigators as described in the their original publication [8]. We limited our analysis to 299 cancer driver genes manually annotated by experts in the pan-cancer field [8]. The MAF*tools* package in the R-statistical environment was used for mutation count calculations within CCS subgroups. A gene was counted as mutated (1) or not (0) for each tumour regardless of the number of mutations within that gene.

### Chromosomal arm-level alterations and Aneuploidy score

Fully processed chromosome arm-level alteration data and tumour aneuploidy scores were accessed from GDC database (https://gdc.cancer.gov) and were derived from Affymetrix SNP 6.0 arrays. All data quality control and processing was performed by the PanCanAtlas investigators as described in the original publication [14]. Chromosome arm-level alterations are presented as estimated ploidy values of +1, 0 and −1 for gains, non-aneuploidy and losses, respectively [9].

### Statistical Analysis

To assess differences among clinico-pathological characteristics of tumour samples and CCS subgroups χ^**2**^ tests were employed. Clinical and survival data were retrieved from the GDC database (https://gdc.cancer.gov/about-data/publications/pancanatlas). Univariate Kaplan-Meier analysis was performed for the CCS in all pan-cancer tumours together and in individual cancer types with Progression Free Interval (PFI) censored at 15 years as the clinical endpoint, as previously recommended [15]. PFI is defined as the period during or after the course of a treatment given to patients in which the disease does not show any progression until a loco-regional recurrence and/or second malignancy occurs, or the patients die from any cause. Multivariable Cox proportional hazard models were used to determine the independent prognostic capacity of the CCS subgroups in all pan-cancer tumours together and in individual cancer types adjusting for cancer type, age (grouped in tertiles), gender, radiation therapy and pathological stage. To compare the prognostic capacity of pan-cancer vs. intra-cancer CCS cutoffs we used the likelihood ratio (LR) which can be interpreted as a goodness-of-fit test. LR and concordance index (c-index) measures were extracted from the output of the coxph function of the *survival* package in R. All statistical analyses were performed using R statistical software version 3.5.3 [16].

## Results

### Cohort clinico-pathological characteristics in relation to CCS subgroups

In line with our aim to compare genomic alterations in tumours with differing levels of cell cycle activity we applied our CCS signature to gene expression data from the tumours of 9,515 pan-cancer patients. Clinico-pathological characteristics for the pan-cancer cohort split by CCS classifications are shown in Table 1. Statistically significant associations were found between patient age, gender, pathological stage, radiotherapy and CCS subgroups (Table 1, Chi-squared test: *P* < 0.001 for all comparisons). After adjusting for cancer type, only stage and radiotherapy remained statistically significant whereby CCS high tumours were more likely to be stage IV and to have received radiotherapy (data not shown).

**Table 1.** Clinical characteristics of all patients split by CCS.

| Variables | Pan-cancer (n = 9515) |  |  | *p* |
| --- | --- | --- | --- | --- |
|  | Low | Intermediate | High |  |
|  | n (%) | n (%) | n (%) |  |
|  | 3145 (33) | 3184 (33.5) | 3186 (33.5) |  |
| Age |  |  |  |  |
| ≤ 54 | 1290 (41) | 876 (28) | 1061 (34) | **< 0.001** |
| 54 - 66 | 1044 (33) | 1062 (33) | 996 (31) |  |
| ≥ 66 | 808 (26) | 1236 (39) | 1119 (35) |  |
| Missing cases = 23 |  |  |  |  |
| Gender |  |  |  |  |
| Male | 1771 (56) | 1372 (43) | 1494 (47) | **< 0.001** |
| Female | 1374 (44) | 1812 (57) | 1692 (53) |  |
| Pathological stage |  |  |  |  |
| Stage I | 859 (45) | 601 (26) | 444 (20) | **< 0.001** |
| Stage II | 480 (25) | 820 (35) | 768 (35) |  |
| Stage III | 419 (22) | 639 (28) | 575 (27) |  |
| Stage IV | 150 (8) | 260 (11) | 382 (18) |  |
| Missing cases & excluded cases<sup>o</sup> = 3118 |  |  |  |  |
| Radiotherapy |  |  |  |  |
| No | 1954 (73) | 2047 (73) | 1820 (65) | **< 0.001** |
| Yes | 709 (27) | 770 (27) | 993 (35) |  |
| Missing cases = 1222 |  |  |  |  |

<sup>o</sup> : I/II NOS-Stage 0/IS/X, , In bold significant $p < 0.05$

### Broad variation in cell cycle activity across cancers and COCA subtypes

We next assessed tumour cell cycle activity by creating pan-cancer, COCA and iCluster boxplots using the continuous CCS. We found the highest levels of cell cycle activity in DLBC, TCGT, HNSC and CESC tumours the lowest in KICH, PCPG, KIRP and PRAD tumours (Figure 1A). Similar results were found using the COCA algorithm - a classification strategy that clusters samples by integrating information from multiple individual cross platform technologies, with CA17 (TCGT) and CA4 (PAN-SCC, mainly HNSC, LUSC and CESC tumours) forming the top two subgroups with the highest cell cycle activity (Figure 1B). CA10 (BRCA, basal-like) and CA25 (Hematologic/lymphatic, mainly THYM and DLBC tumours), also showed high cell cycle activity, whilst CA1 (CNS/Endocrine, mainly PCPG tumours), CA14 (PRAD) and C21 (PAN-Kidney) showed the lowest levels of all COCA subtypes (Figure 1B). Analogous results were noted using the iCluster classification strategy (Supplemental Figure 2). Examining cell cycle activity clusters using heatmap analysis demonstrated that tumours with low levels of cell cycle activity (and thus classified as CCS Low) show low expression of the majority of genes in all cell cycle phases (G1 to M), whilst the opposite is true for tumours with high levels of cell cycle activity (Figure 1C, compare tumours with black column-side colour to those with yellow).

**Figure 1.**
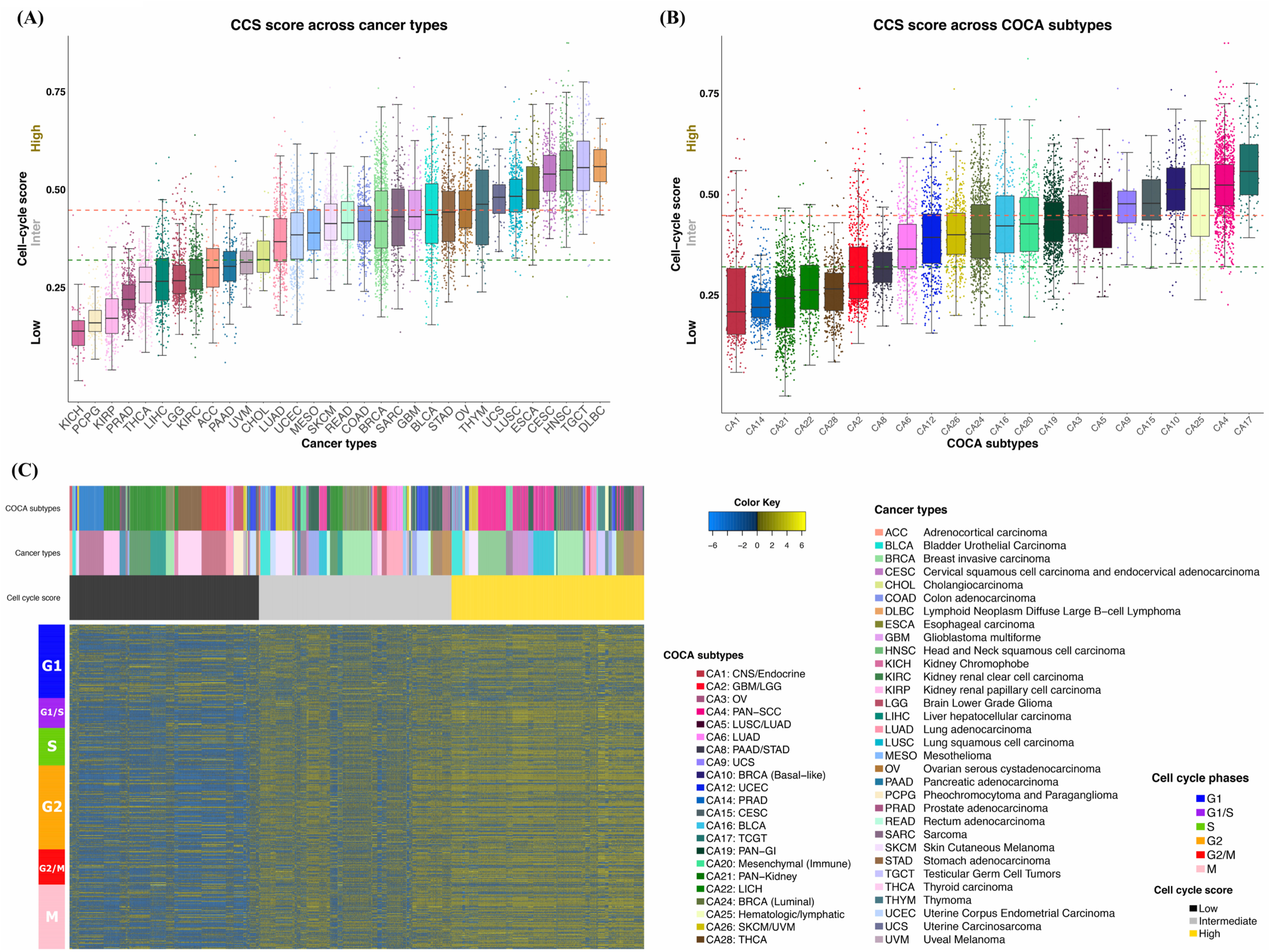
CCS score across cancer types and COCA subgroups. Boxplots comparing CCS across (A) pan-cancer types and (B) COCA subtypes. (C) Heatmap of CCS genes across pan-cancer tumours. Heatmap colside colours (horizontal, above heatmap) represent cell cycle score, cancer types and COCA groups as indicated in figure legend. Rowside colours (vertical, left hand side of heatmap) represent cell cycle phases.

### *TP53* and *PIK3CA* mutations display increasing frequency across cell cycle activity subgroups

To more clearly delineate the frequency of DNA mutations in relation to the magnitude of cell cycle activity we next examined the mutational frequency of 299 well defined oncogene and tumour suppressor driver genes within CCS subgroups. *TP53* was found to be the most mutated gene in all three CCS subgroups and displayed an increase in mutational frequency with increasing CCS activity (Figure 2A, Supplemental Table 1, Chi-squared test: *P* < 0.001). In CCS Low tumours 40% of *TP53* mutations were found in LGG, whereas in CCS high tumours *TP53* mutations were most common in HNSC (18%), LUSC (17%) and BRCA (13%) (Highlighted in Figure 2A). *PIK3CA* was the second-most commonly mutated gene in CCS intermediate and high tumours and fifth most common in CCS Low tumours (Figure 2A). It is also more frequently mutated in CCS Intermediate and High tumours relative to CCS Low (Supplemental Table 1, *P* < 0.001). *PIK3CA* mutations in BRCA and UCEC were common across all CCS subgroups and were additionally found in HNSC and CESC in CCS high tumours (Figure 2A). Of interest, whilst *BRAF* mutations were prominent in Low and Intermediate subgroups as the third and eleventh most mutated gene respectively, it was absent from the top 15 in CCS High tumours (Figure 2A, red arrows, Supplemental Table 1, *P* = 0.001). This suggests that other genes are more commonly mutated in tumours with high cell cycle activity. The top 50 most frequently mutated genes in all CCS subgroups are shown in Supplemental Table 2.

**Figure 2.**
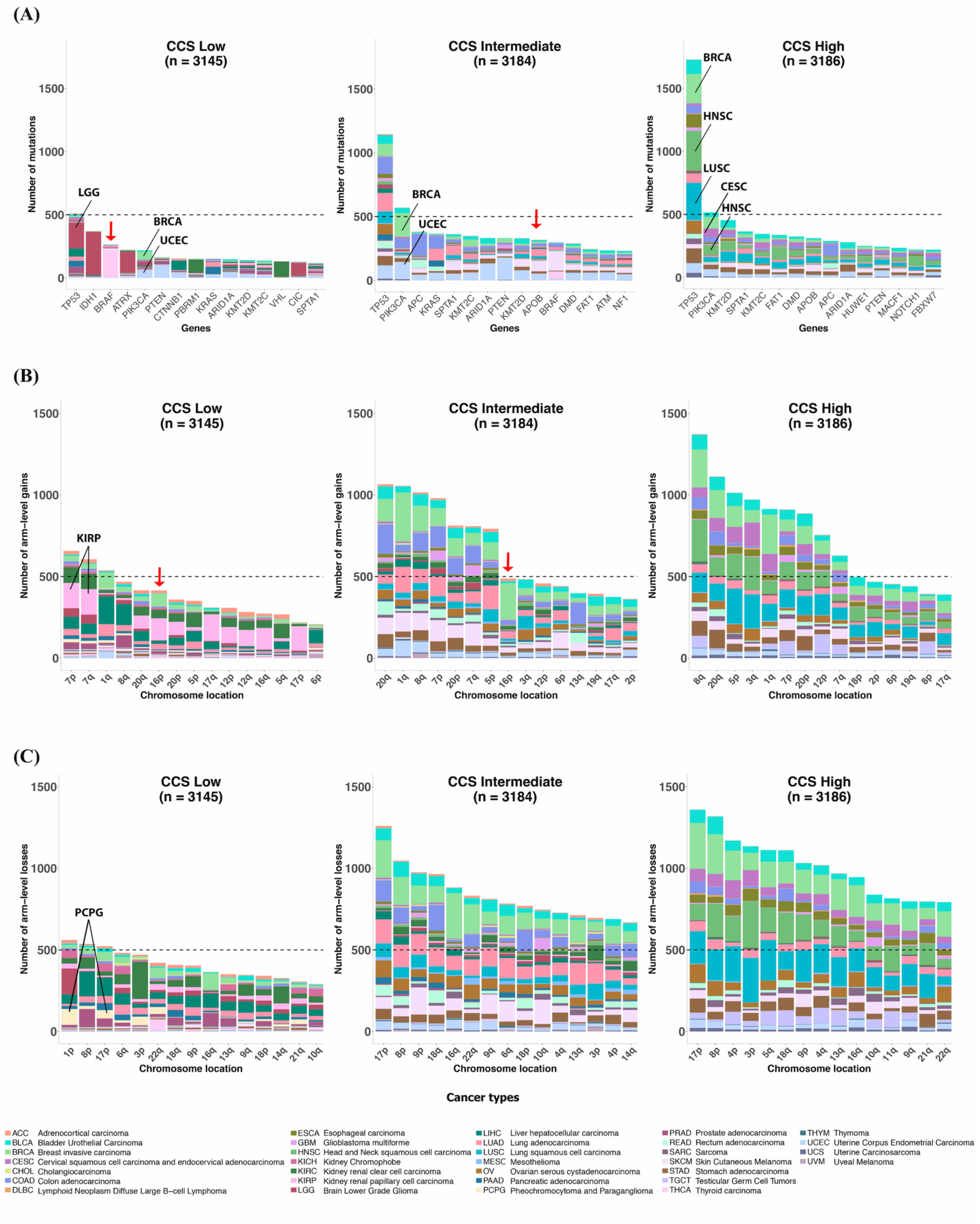
Top 15 most commonly mutated genes or chromosomal arm-level alterations within CCS subgroups. Pan-cancer tumours were divided into tertiles on the basis of low, intermediate or high CCS. Within each subgroup the Top 15 (A) Most frequently mutated oncogenes and tumour suppressor genes, (B) Arm-level gains and (C) Arm-level losses are shown. Cancer type colour key is are shown at the bottom of the figure. Red arrows indicate *BRAF* mutations and 16p gains in CCS low and intermediate subgroups.

### Higher levels of chromosomal gains and losses in CCS intermediate and high tumours

We next performed the same subgroup analysis, but this time focusing on chromosome arm-level gains and losses. All CCS subgroups showed a high number of gains to arms 20q, 8q and 7p and losses to arms 17p and 8p (Figure 2B and C, respectively, all CCS subgroups). Moreover, these chromosomal aberrations all displayed an increase in frequency with increasing CCS activity (Supplemental Table 1, Chi-squared test: *P* < 0.001 for all comparisons, not adjusted for multiple testing). Overall, gains in KIRP (Figure 2B, highlighted) and losses in PCPG cancers (Figure 2C, highlighted) were more common CCS Low tumours relative to CCS Intermediate and High subgroups, as could be anticipated given the low cell cycle activity levels displayed by these tumour types and their grouping into the CCS Low tumour subgroup (Figure 1A). Analogous to our *BRAF* mutation findings, gains to 16p (Figure 2B, red arrows) were more common in CCS Low and Intermediate subgroups relative to the CCS High subgroup (Supplemental Table 1, *P* < 0.001). The frequency of chromosomal arm gains and losses in all CCS subgroups are shown in Supplemental Table 3.

Next, we examined genomic alterations more broadly within CCS subgroups and found the frequency of gene mutations and chromosomal arm gains and losses to be greater in CCS Intermediate and High groups relative to Low (Figure 3 A - C, Tukey HSD test, 3A top 50 DNA mutations: *P* < 0.001 and *P* < 0.001, 3B chromosomal gains: *P* = 0.018 and *P* < 0.001 and losses 3C: *P* < 0.001 and *P* < 0.001 for Low vs. Intermediate and High, respectively). Similarly, using the recently derived aneuploidy score [9] – a measure of the total number of chromosome arms with arm-level copy number changes in a given sample, we also found a statistically significant increase with increasing CCS activity levels (Figure 3 D, *P* < 0.001 for all comparisons).

**Figure 3.**
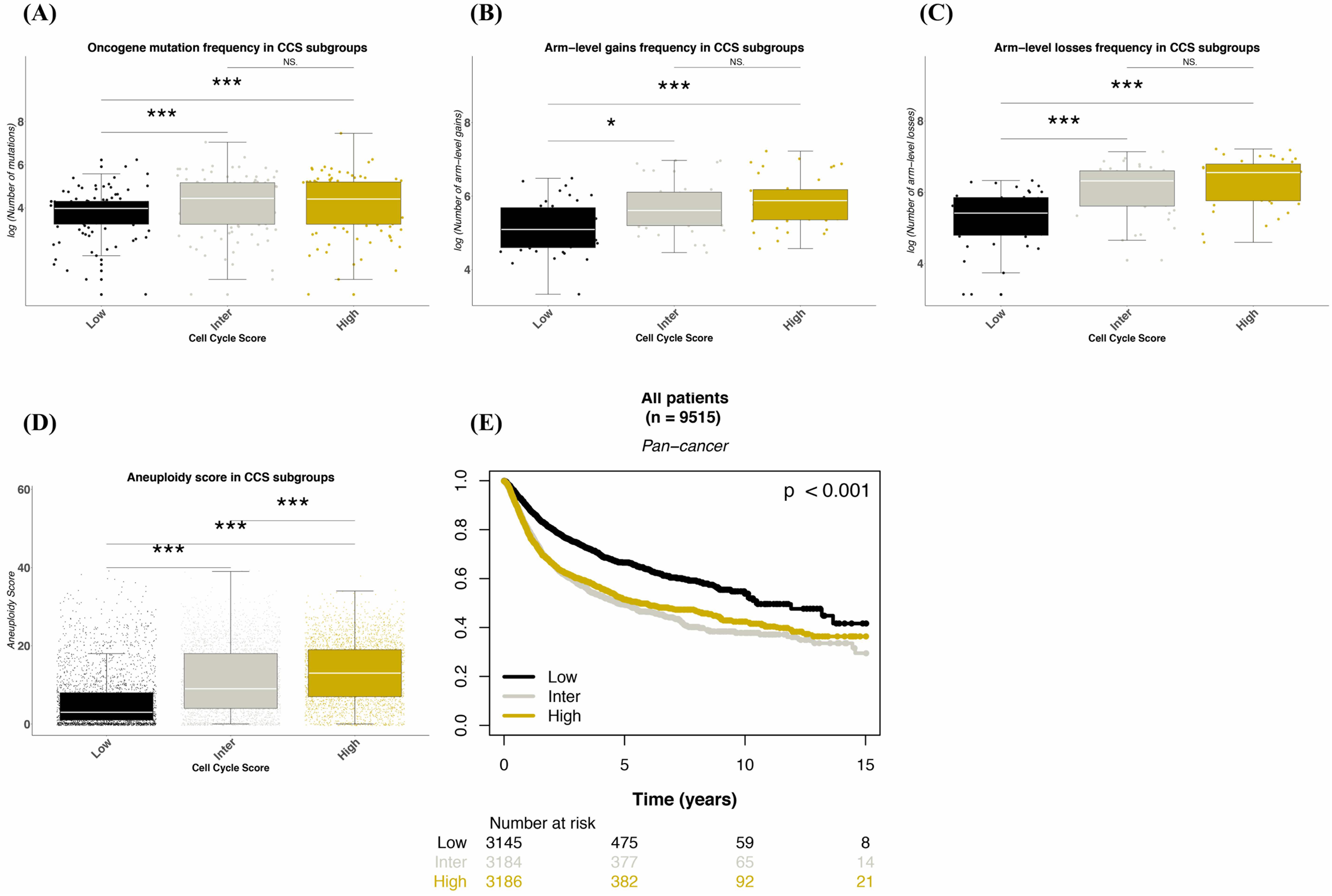
Boxplots comparing frequency of DNA alterations across CCS subgroups. Pan-cancer tumours were divided into tertiles on the basis of low, intermediate or high CCS. Within each subgroup the number of (A) Total mutations in the top 50 most mutated oncogenes or tumour suppressor genes, (B) Total chromosomal arm-level gains, (C) Total chromosomal arm-level losses and (D) Aneuploidy score are shown. (E) Kaplan-Meier analysis of CCS subgroups with Progression-free Interval (PFI) as clinical endpoint. Low/Inter/High = Low/Intermediated/High CCS subgroups, *p* values in boxplots (based on ANOVA with post-hoc Tukey HSD test) = NS > 0.05, * < 0.05, ** < 0.01, *** < 0.001; *p* value in the Kaplan-Meier curves refer to long-rank tests.

### CCS signature provides independent prognostic information at pan-cancer level

We next assessed the relationship between CCS and PFI using Kaplan-Meier and multivariable Cox proportional hazard regression model analyses. In univariate Kaplan-Meier analysis patients whose tumours were classified as CCS Low had a significantly longer PFI relative to those classified as CCS Intermediate or High (Figure 3 E, log-rank test: *P* < 0.001). This significance remained when adjusting for tumour type, age, gender, pathological stage and radiotherapy in Cox proportional hazard analysis (Table 2, CCS intermediate: HR 1.37 95% CI 1.17 – 1.60, CCS high: HR 1.54 95% CI 1.29 – 1.84, tumour type not shown). The upper Age tertile (≥ 66) remained statistically significant in the same model (HR 1.19 95% CI 1.05 – 1.35 vs. Ref), as did all pathological stages vs. the Stage I model reference group.

**Table 2.** Multivariable Cox-regression analysis of the CCS signature across pan-cancer patients.

| Variables | Pan-cancer (n = 5421)* |  |  |  |
| --- | --- | --- | --- | --- |
|  | N (%) | HR | 95% CI | *p* |
| Age |  |  |  |  |
| ≤ 54 | 1679 (31) | Ref | - | - |
| 54 - 66 | 1761 (32) | 1.04 | 0.91 - 1.18 | 0.551 |
| ≥ 66 | 1981 (37) | 1.19 | 1.05 - 1.35 | **0.008** |
| Missing cases = 23 |  |  |  |  |
| Gender |  |  |  |  |
| Male | 2757 (51) | Ref | - | - |
| Female | 2664 (49) | 0.96 | 0.87 - 1.07 | 0.483 |
| Pathological stage |  |  |  |  |
| Stage I | 1561 (29) | Ref | - | - |
| Stage II | 1852 (34) | 1.60 | 1.38 - 1.86 | **< 0.001** |
| Stage III | 1378 (25) | 2.41 | 2.08 - 2.79 | **< 0.001** |
| Stage IV | 630 (12) | 5.04 | 4.21 - 6.03 | **< 0.001** |
| Missing cases = 3118 |  |  |  |  |
| Radiotherapy |  |  |  |  |
| No | 3997 (74) | Ref | - | - |
| Yes | 1424 (26) | 0.97 | 0.84 - 1.11 | 0.658 |
| Missing cases = 1222 |  |  |  |  |
| Cell cycle score |  |  |  |  |
| Low | 1505 (28) | Ref | - | - |
| Intermediate | 2013 (37) | 1.37 | 1.17 - 1.60 | **< 0.001** |
| High | 1903 (35) | 1.54 | 1.29 - 1.84 | **< 0.001** |

\*: adjusted for cancer types, Ref: Reference groups, N: Number of patients, HR: hazard ratio, CI: confidence interval, In bold significant $p < 0.05$

In order to determine if the prognostic capacity of the CCS was similar in all cancer types, we again performed Kaplan-Meier and Cox proportional hazard modelling but this time focusing on individual cancers. CCS provided significant independent prognostic information in seven cancer types: KIRC (*P* = 0.006), LGG (*P* < 0.001), LUAD (*P* = 0.031), PAAD (*P* = 0.026), SARC (*P* < 0.001), UCEC (*P* = 0.012) and UVM (*P* = 0.001, Supplemental Figure 3, alphabetical ordering, unadjusted for multiple testing). Finally, as the CCS subgroups are based on a tertile split of cell cycle activity on a pan-cancer level, we hypothesised that deriving subgroups in this manner may provide superior prognostic information to a simple tertile split within (intra) each cancer type. To test this hypothesis, we compared our pan-cancer CCS tertile subgroups to intra-cancer CCS tertile subgroups. We found that whilst both cut-offs provide significant prognostic information in the above seven cancer types (Compare Kaplan-Meier curves for pan-cancer CCS to intra-cancer CCS, Supplemental Figure 4), a pan-cancer cut-off provides more prognostic information in KIRC (LR = 24.7), LGG (LR = 31.1), SARC (LR = 18.5) and UVM cancers (LR = 17.1, Table 3, compare pan-cancer column to intra-cancer). These findings suggest that deriving transcriptional biomarker cut-points on a pan-cancer level may be advantageous relative to deriving them in a single cancer type.

**Table 3.** Comparison of the prognostic value of *pan-cancer* vs *intra-cancer* cutoffs for the CCS signature.

| Models | Cell cycle score |  |  |  |  |  |  |  |
| --- | --- | --- | --- | --- | --- | --- | --- | --- |
|  | Total | Events | *Pan-Cancer* |  |  | *Intra-cancer* |  |  |
| | | | C-index | LR-$X^2$ | *p* | C-index | LR-$X^2$ | *p* |
| Univariate |  |  |  |  |  |  |  |  |
| KIRC | 480 | 148 | 0.589 | 24.7 | < **0.001** | 0.567 | 9.9 | **0.002** |
| LGG | 507 | 192 | 0.609 | 31.1 | < **0.001** | 0.627 | 21.9 | < **0.001** |
| LUAD | 498 | 188 | 0.556 | 6.6 | **0.010** | 0.560 | 7.7 | **0.006** |
| PAAD | 158 | 90 | 0.588 | 9.7 | **0.002** | 0.611 | 13.7 | < **0.001** |
| SARC | 242 | 124 | 0.610 | 18.5 | < **0.001** | 0.614 | 14.3 | < **0.001** |
| UCEC | 514 | 105 | 0.555 | 5.5 | **0.019** | 0.589 | 9.9 | **0.002** |
| UVM | 80 | 24 | 0.732 | 17.1 | < **0.001** | 0.707 | 11.5 | < **0.001** |

KIRC: Kidney renal clear cell carcinoma; LGG: Brain Lower Grade Glioma; LUAD: Lung adenocarcinoma; PAAD: Pancreatic adenocarcinoma; SARC: Sarcoma;
UCEC: Uterine Corpus Endometrial Carcinoma; UVM: Uveal Melanoma; Events: Number of deaths from each cancer type; CCS: Cell cycle score; LR- $X^2$ : Likelihood ratio;C-index: Concordance index; In bold significant $p < 0.05$

## Discussion

The present study integrates gene expression, DNA mutation, DNA copy number and clinico-pathological data from 9,515 pan-cancer patients in order to better understand the DNA level alterations present in tumours with low, intermediate and high cell cycle activity. Our main findings show first, that cell cycle activity varies broadly across and within cancer types; second, that *TP53, PIK3CA* and chromosomal alterations (including gains to 20q, 8q, 7p and losses to arms 17p and 8p) occur with increasing frequency in tumours with increasing cell cycle activity; third, whilst in general similar mutations/arm level alterations are present within tumours with low, intermediate and high cell cycle activity, mutations in *BRAF*, gains in 16p and losses in 6q were less frequent in tumours with high cell cycle activity; and fourth, that deriving cut-points for biomarkers on a pan-cancer level may provide more prognostic information than deriving them within specific cancer types. These analyses are the first to provide broad insight on the genetic alterations occurring within tumours grouped on the basis of cell cycle activity in order to advance our understanding of a pathway that is frequently dysregulated in human malignancies.

In pan-cancer analyses, *TP53, PIK3CA, KRAS, PTEN* and *ARID1A* genes have all been previously demonstrated to be mutated in over 15 different cancer types [8]. These genes also featured heavily in our mutational analysis with *TP53* and *PIK3CA* mutations showing the high mutational frequency across CCS subgroups. This implies that mutations in these genes are found in tumours with a broad range of cell cycle activity and are not just associated with highly cycling cancers, despite their very clear links to cell cycle progression [17,18]. Whilst we found the *ARID1A* gene to be mutated in all CCS subgroups *BRAF* was notable for only being found in the top 15 of the CCS Low and Intermediate subgroups, implying that other genes are more commonly mutated in tumours with high cell cycle activity, such as *TP53* and *PIK3CA*.

It has recently been demonstrated that tumour aneuploidy is inversely correlated to immune signalling genes and positively correlated to cell cycle and pro-proliferation pathways [9]. Our findings are in line with these showing a step wise increase in aneuploidy score with increasing CCS activity levels. Related to this, whilst most of predominant chromosome arm-level alterations we observed overlapped with those from the pan-cancer publication [9], our within subgroup analysis yielded some novel findings. In particular, and analogous to our mutational results, we found that specific gains (16p) and losses (6q) were present in the CCS Intermediate and high subgroups only (Figure 2B and C, red arrows). This raises the possibility that these chromosomal alterations could potentially be used as novel clinical biomarkers for more indolent tumours on a pan-cancer level.

We found that our cell cycle score gene expression signature, previously established in a breast cancer setting [11,12], provided independent prognostic information on a pan-cancer level. This signature was originally conceived as simple biological measure of cell cycle activity in response to the dependence of more established commercial gene expression signatures on multiple cell cycle/cell proliferation genes for their prognostic capacity [19]. In keeping with its descriptive nature, we have not attempted to maximise the signature’s prognostic capacity through selection of genes that are the strongest predictors of the study’s clinical endpoint - progression free interval. Despite this, the signature performed well in both Kaplan-Meier and multi-variable analyses, likely owing to its ability to select for faster growing, more aggressive tumours. Following on from these results we also noted that deriving CCS tertiles of activity on a pan-cancer level may provide more prognostic information than deriving them within a specific cancer type. This may be of utility in a clinical setting where a gene transcript is being used as a biomarker for treatment response, such as the recent example of cyclin E expression and Palbociclib efficacy in metastatic breast cancer patients [20]. In this instance it is conceivable that re-defining a cyclin E cut-point on the basis of pan-cancer expression levels of the gene may more clearly delineate which patients are likely to be resistant to the drug.

There are three main strengths to our study; first, we utilise a novel methodology to examine the DNA alterations in subgroups of tumours that is based on the magnitude of cell cycle activity both across and within cancer types; second, our analysis provides an expansive overview of the frequency of DNA mutations and chromosomal gains and losses in subgroups of low, intermediate and high cell cycle activity; and third, we demonstrate the translational relevance of our work by relating our CCS signature to a clinical survival endpoint – PFI. The limitations are as follows; first, our analysis focuses on DNA and RNA technologies only, with no protein or methylation array data included; second, we chose to study broad chromosomal gains and losses rather than gene-centric copy number changes – this was to avoid a situation where the most changed genes within a given CCS subgroup would all come from the same chromosomal location; and third, no external validation was performed for the CCS signature, although we are not aware of any other pan-cancer dataset where it could be validated and more importantly, we are not currently proposing it for use in a clinical setting – rather as a general tool to examine the cell cycle activity of a given tumour.

In summary, this study describes the DNA mutations and chromosomal alterations found in tumours with low, intermediate and high levels of cell cycle activity and also demonstrates the ability of a simple cell cycle gene expression signature to provide independent prognostic information at a pan-cancer level.

## Abbreviations

ACC: Adrenocortical carcinoma
BLCA: Bladder Urothelial Carcinoma
CCS: Cell Cycle Score
CDKs: Cyclin dependent kinases
CESC: Cervical squamous cell carcinoma and endocervical adenocarcinoma
CHOL: Cholangiocarcinoma
c-index: Concordance index
COAD: Colon adenocarcinoma
COCA: Cluster of cluster
ESCA: Esophageal carcinoma
DLBC: Lymphoid Neoplasm Diffuse Large B-cell Lymphoma
GBM: Glioblatoma multiforme
GDC: Genomic data commons
HNSC: Head and Neck squamous cell carcinoma
iCluster: Integrative Cluster
KICH: Kidney Chromophobe
KIRC: Kidney renal clear cell carcinoma
KIRP: Kidney renal papillary cell carcinoma
LGG: Brain lower grade Glioma
LICH: Liver hepatocellular carcinoma
LR: Likelihood ratio
LUAD: Lung adenocarcinoma
LUSC: Lung squamous cell carcinoma
NIH: National Institutes of Health
OV: Ovarian serous cystadenocarcinoma
PAAD: Pancreatic adenocarcinoma
PCPG: Pheochromocytoma and Paraganglioma
PFI: Progression Free Interval
PRAD: Prostate adenocarcinoma
READ: Rectum adenocarcinoma
SARC: Sarcoma
SKCM: Skin Cutaneous Melanoma
STAD: Stomach adenocarcinoma
TCGA: The Cancer Genome Atlas
TGCT: Testicular Germ Cell tumours
THYM: Thymoma
THCA: Thyroid carcinoma
UCS: Uterine Carcinosarcoma
UCEC: Uterine Corpus Endometrial Carcinoma
UVM: Uveal Melanoma.

## Availability of data and materials

The data used in this study are publicly available on NIH (https://gdc.cancer.gov/about-data/publications/pancanatlas).

## Authors’ contributions

A.L. and N.P.T contributed the study concept and design. A.L. contributed to the acquisition and analysis of data. All authors interpreted the data and did the manuscript drafting and critical revision. N.P.T did the study supervision. All authors read and approved the final manuscript.

