## Supplementary material for "A pan-cancer analysis of the frequency of DNA alterations across cell cycle activity levels": Suplemental Figures and Tables

**Supplemental figure 1.**

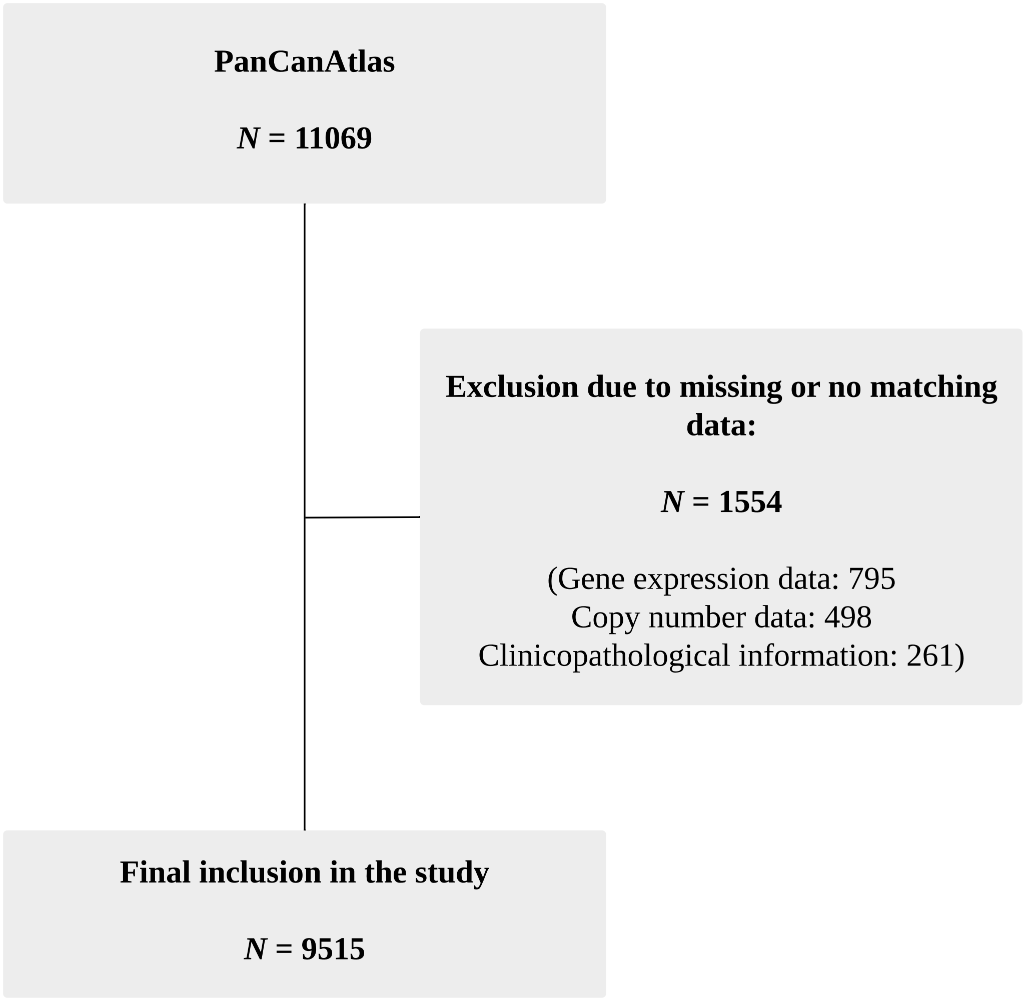


**Supplemental figure 1. CONSORT diagram of patient selection**

PanCanAtlas = The Pan-Cancer Atlas project including 32 cancer types

### Supplemental figure 2.

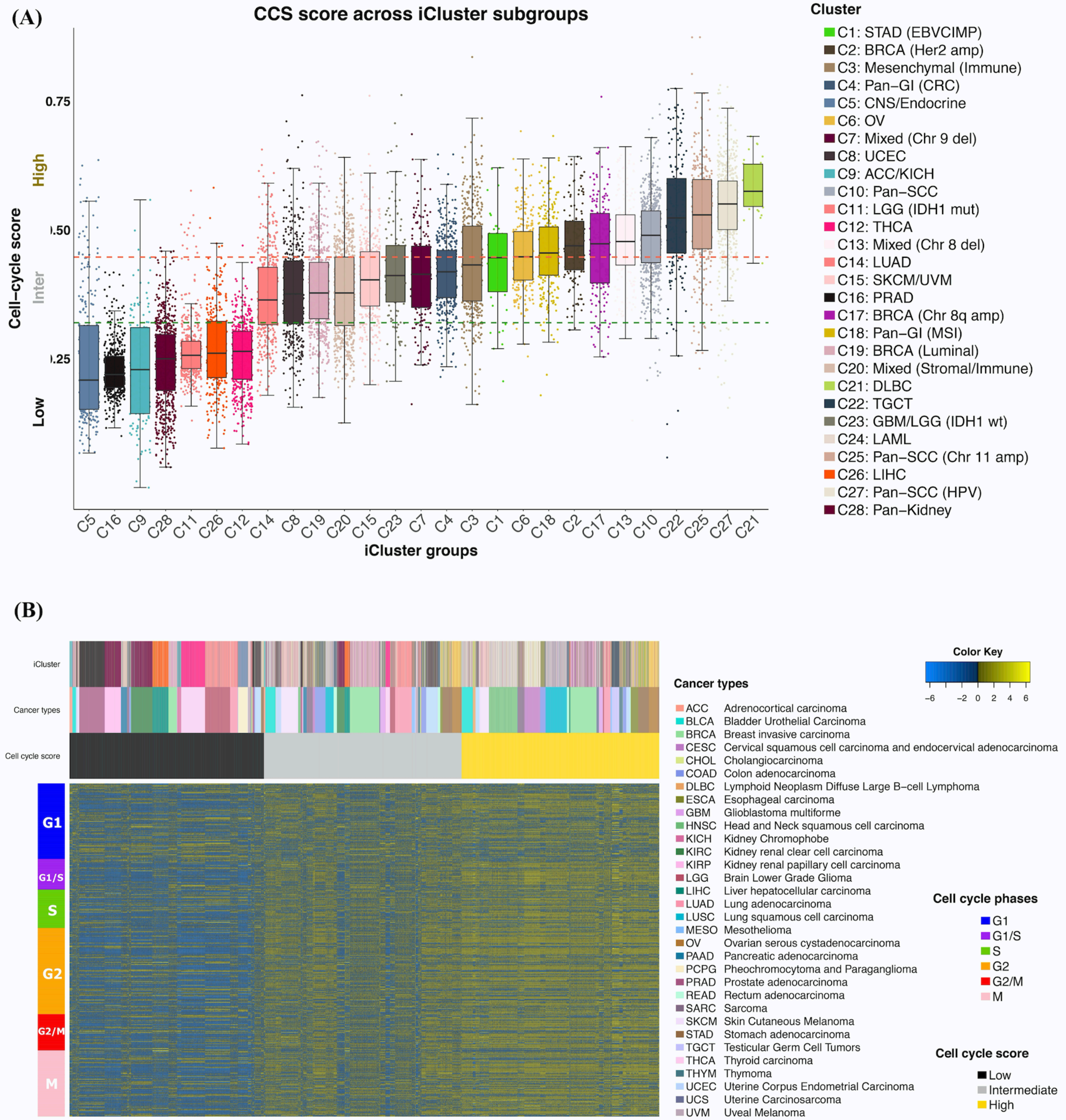


**Supplemental figure 2. Pan-cancer cell cycle score (CCS) in integrative cluster (iCluster) subgroups**

(A) Boxplots comparing CCS across iCluster subgroups and (B) Heatmap of CCS genes across pan-cancer patients. Heatmap colside colours (horizontal) represent cell cycle score, cancer types and iCluster groups. Rowside colours represent cell cycle phases.

### Supplemental figure 3.

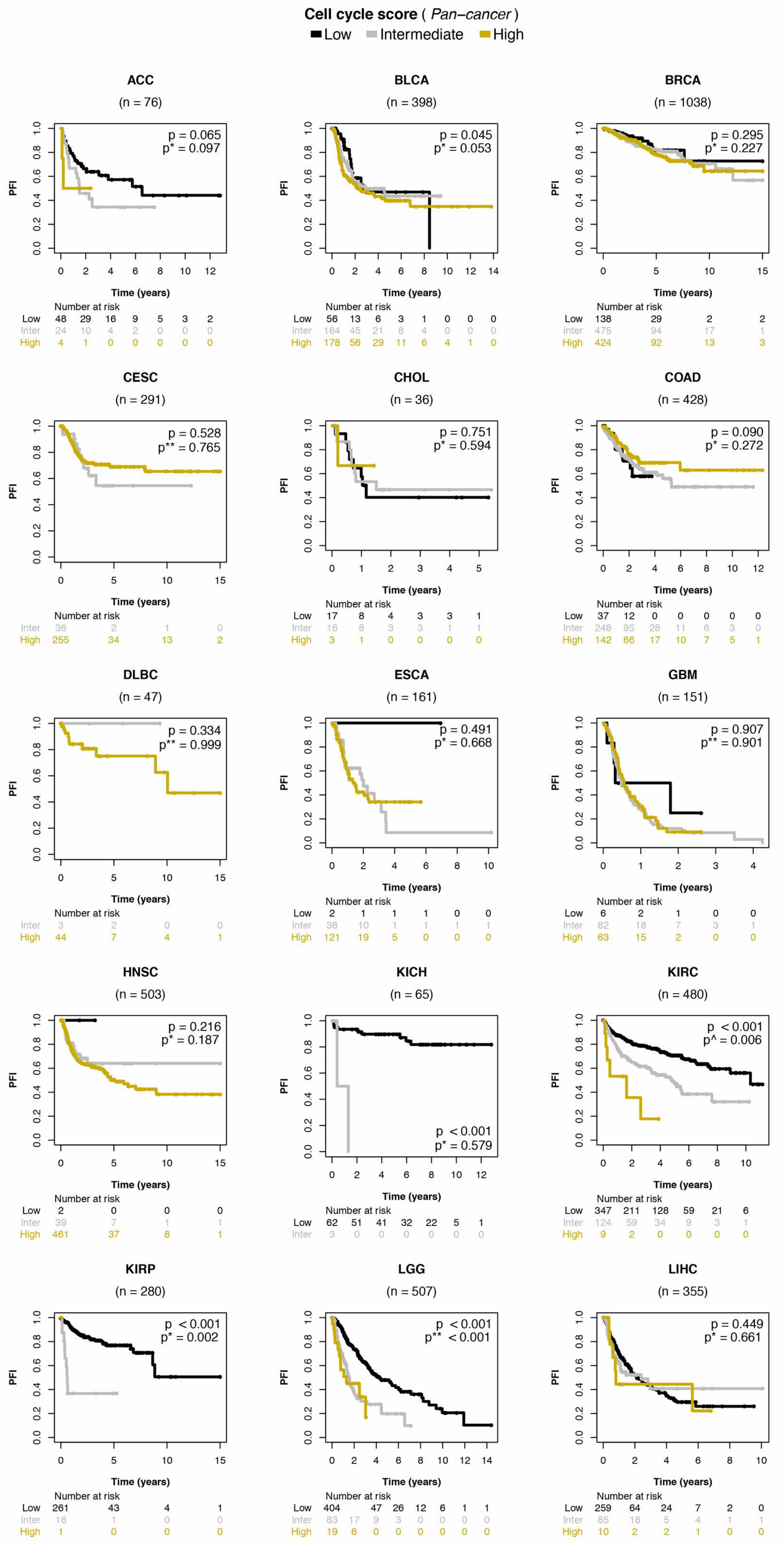

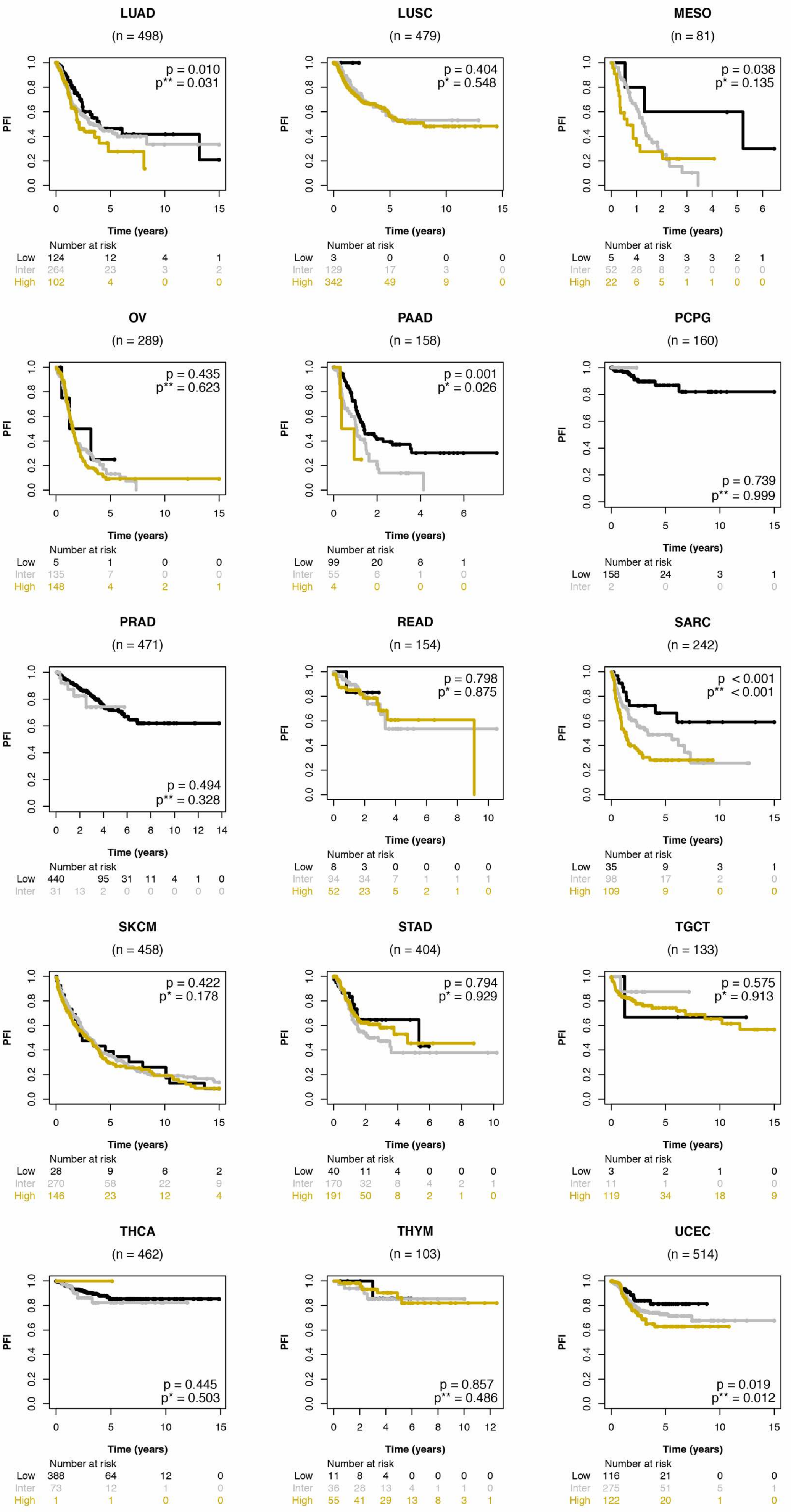
LUAD
(n = 498)
p = 0.010
p** = 0.031
PFI
Time (years)
Number at risk
Low 124 12 4 1
Inter 264 23 3 2
High 102 4 0 0
LUSC
(n = 479)
p = 0.404
p* = 0.548
Number at risk
Low 3 0 0 0
Inter 129 17 3 0
High 342 49 9 0
MESO
(n = 81)
p = 0.038
p* = 0.135
Number at risk
Low 5 4 3 3 3 2 1
Inter 52 28 8 2 0 0 0
High 22 6 5 1 1 0 0
OV
(n = 289)
p = 0.435
p** = 0.623
Number at risk
Low 5 1 0 0
Inter 135 7 0 0
High 148 4 2 1
PAAD
(n = 158)
p = 0.001
p* = 0.026
Number at risk
Low 99 20 8 1
Inter 55 6 1 0
High 4 0 0 0
PCPG
(n = 160)
p = 0.739
p** = 0.999
Number at risk
Low 158 24 3 1
Inter 2 0 0 0
PRAD
(n = 471)
p = 0.494
p** = 0.328
Number at risk
Low 440 95 31 11 4 1 0
Inter 31 13 2 0 0 0 0 0
READ
(n = 154)
p = 0.798
p* = 0.875
Number at risk
Low 8 3 0 0 0 0
Inter 94 34 7 1 1 1
High 52 23 5 2 1 0
SARC
(n = 242)
p < 0.001
p** < 0.001
Number at risk
Low 35 9 3 1
Inter 98 17 2 0
High 109 9 0 0
SKCM
(n = 458)
p = 0.422
p* = 0.178
Number at risk
Low 28 9 6 2
Inter 270 58 22 9
High 146 23 12 4
STAD
(n = 404)
p = 0.794
p* = 0.929
Number at risk
Low 40 11 4 0 0 0
Inter 170 32 8 4 2 1
High 191 50 8 2 1 0
TGCT
(n = 133)
p = 0.575
p* = 0.913
Number at risk
Low 3 2 1 0
Inter 11 1 0 0
High 119 34 18 9
THCA
(n = 462)
p = 0.445
p* = 0.503
Number at risk
Low 388 64 12 0
Inter 73 12 1 0
High 1 1 0 0
THYM
(n = 103)
p = 0.857
p** = 0.486
Number at risk
Low 11 8 4 0 0 0 0
Inter 36 28 13 4 1 1 0
High 55 41 29 13 8 3 1
UCEC
(n = 514)
p = 0.019
p** = 0.012
Number at risk
Low 116 21 0 0
Inter 275 51 5 1
High 122 20 1 0

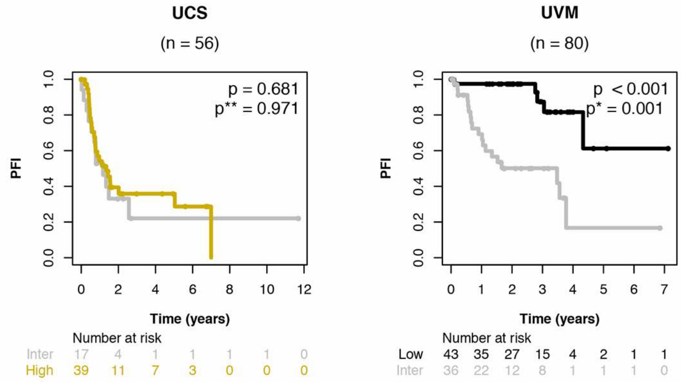


**Supplemental figure 3. Kaplan-Meier analysis of CCS within individual cancer types, Progression-free Interval (PFI) as clinical endpoint**

ACC: Adrenocortical carcinoma, BCLA: Bladder Urothelial Carcinoma, BRCA: Breast invasive carcinoma, CESC: Cervical squamous cell carcinoma and endocervical adenocarcinoma, CHOL: Cholangiocarcinoma, COAD: Colon adenocarcinoma, DLBC: Lymphoid Neoplasm Diffuse Large B-cell Lymphoma, ESCA: Esophageal carcinoma, GBM: Glioblastoma multiforme, HNSC: Head and Neck squamous cell carcinoma, KICH: Kidney Chromophobe, KIRC: Kidney renal clear cell carcinoma, KIRP: Kidney renal papillary cell carcinoma, LGG: Brain Lower Grade Glioma, LIHC: Liver hepatocellular carcinoma, LUAD: Lung adenocarcinoma,  LUSC: Lung squamous cell carcinoma, MESO: Mesothelioma, OV: Ovarian serous cystadenocarcinoma, PAAD: Pancreatic adenocarcinoma , PCPG: Pheochromocytoma and Paraganglioma, PRAD: Prostate adenocarcinoma, READ: Rectum adenocarcinoma, SARC: Sarcoma, SKCM: Skin Cutaneous Melanoma, STAD: Stomach adenocarcinoma, TGCT: Testicular Germ Cell Tumors, THCA: Thyroid carcinoma, THYM: Thymoma, UCEC: Uterine Corpus Endometrial Carcinoma, UCS: Uterine Carcinosarcoma, UVM: Uveal Melanoma;
Low/Inter/High: Low/Intermediate and High CCS subgroups; *p* value refer to log-rank tests; *p** : adjusted *p* value for age, gender, radiation therapy and pathalogical stage; *p*** : adjusted *p* value for age, gender, radiation therapy, *p^* : adjusted *p* value for age, gender, pathological stage

### Supplemental figure 4.

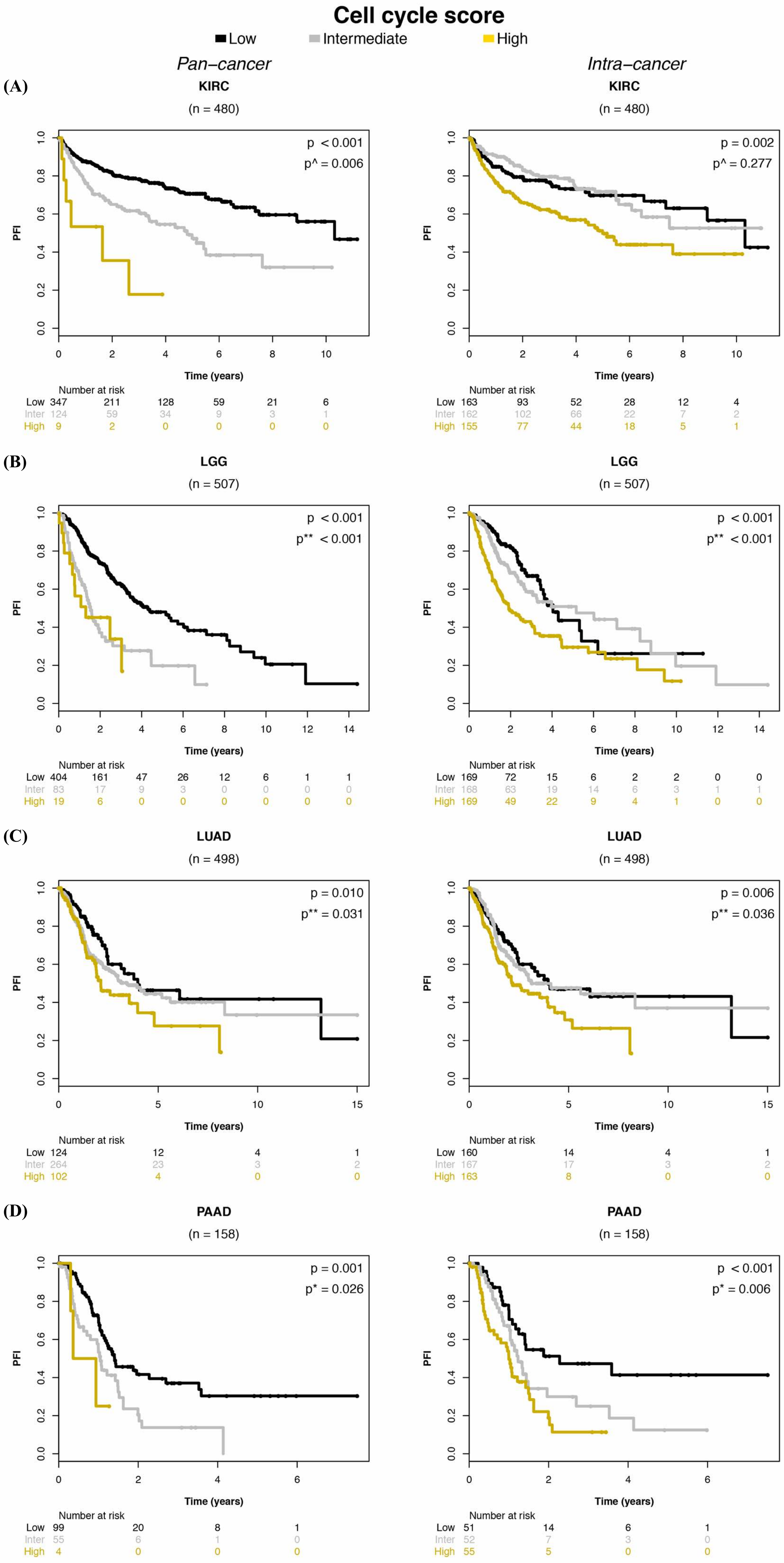

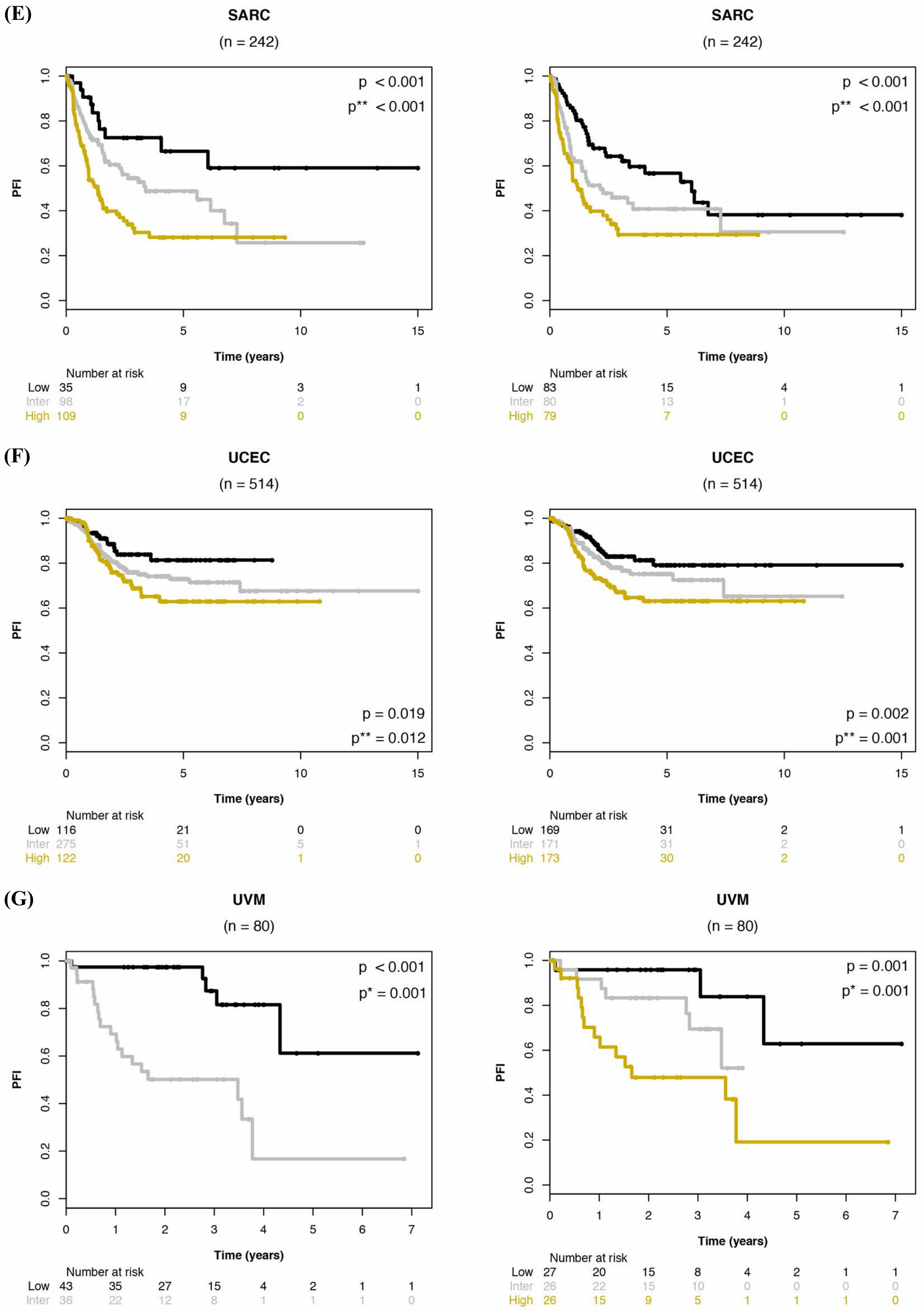


**Supplemental figure 4. Kaplan-Meier analysis of CCS comparing pan-cancer versus intra-cancer cutpoints, Progression-free Interval (PFI) as clinical endpoint.**

Pan-cancer (left hand side) and Intra-cancer (right hand side) survival curves for (A) KIRC: Kidney renal clear cell carcinoma, (B) LGG: Brain lower grade glioma, (C) LUAD: Lung adenocarcinoma, (D) PAAD: Pancreatic adenocarcinoma, (E) SARC: Sarcoma, (F) UCEC: Uterine corpus endometrial carcinoma and (G) UVM: Uveal melanoma. Low/Inter/High; Low/Inter/High: Low/Intermediate and High CCS subgroups; *p* value refer to log-rank tests; *p** : adjusted *p* value for age, gender, radiation therapy and pathalogical stage; *p*** : adjusted *p* value for age, gender, radiation therapy, *p*^ : adjusted *p* value for age, gender, pathological stage

#### Supplemental Table 1

**Number of mutated genes & chromosomal arm-level gains/losses vs non-mutated cases split by Cell cycle score**

| Variables | | Cell cycle score Low n (%) | Intermediate n (%) | High n (%) | *p* |
|---|---|---|---|---|---|
| | | 3145 (100) | 3184 (100) | 3186 (100) | |
| **Genes** | | | | | |
| TP53 | Mutated | 510 (16) | 1146 (36) | 1729 (54) | **< 0.001** |
| | Non-mutated | 2635 (84) | 2038 (64) | 1457 (46) | |
| PIK3CA | Mutated | 219 (7) | 570 (18) | 517 (16) | **< 0.001** |
| | Non-mutated | 2926 (93) | 2614 (82) | 2669 (84) | |
| BRAF | Mutated | 265 (8) | 298 (9) | 152 (5) | **< 0.001** |
| | Non-mutated | 2880 (92) | 2886 (91) | 3034 (95) | |
| **Chromosome arm-level gains** | | | | | |
| 20q | Gain | 416 (14) | 1066 (37) | 1113 (39) | **< 0.001** |
| | No gain | 2663 (87) | 1844 (63) | 1725 (61) | |
| | | missing cases = 67 | missing cases = 274 | missing cases = 348 | |
| 8q | Gain | 468 (16) | 1016 (37) | 1373 (53) | **< 0.001** |
| | No gain | 2475 (84) | 1714 (63) | 1194 (47) | |
| | | missing cases = 202 | missing cases = 454 | missing cases = 619 | |
| 7p | Gain | 657 (21) | 980 (33) | 911 (32) | **< 0.001** |
| | No gain | 2429 (79) | 1990 (67) | 1889 (68) | |
| | | missing cases = 59 | missing cases = 214 | missing cases = 386 | |
| 16p | Gain | 415 (15) | 488 (17) | 372 (13) | **< 0.001** |
| | No gain | 2677 (85) | 2425 (83) | 2396 (87) | |
| | | missing cases = 53 | missing cases = 271 | missing cases = 418 | |
| **Chromosome arm-level losses** | | | | | |
| 17p | Loss | 523 (17) | 1260 (43) | 1360 (47) | **< 0.001** |
| | No loss | 2467 (83) | 1673 (57) | 1505 (53) | |
| | | missing cases = 155 | missing cases = 251 | missing cases = 321 | |
| 8p | Loss | 537 (18) | 1047 (36) | 1319 (47) | **< 0.001** |
| | No loss | 2388 (82) | 1827 (64) | 1475 (53) | |
| | | missing cases = 220 | missing cases = 310 | missing cases = 392 | |

Correlations were calculated using $\chi 2$ test, In bold significant $p < 0.05$

#### Supplemental Table 2

**Top 50 Genes with highest number of mutations within each Cell Cycle score subgroup**

| Rank | Cell Cycle Score | | | | | |
|---|---|---|---|---|---|---|
| | Low | | Intermediate | | High | |
| | Genes | Total # | Genes | Total # | Genes | Total # |
| 1 | TP53 | 510 | TP53 | 1146 | TP53 | 1729 |
| 2 | IDH1 | 372 | PIK3CA | 570 | PIK3CA | 517 |
| 3 | BRAF | 265 | APC | 381 | KMT2D | 455 |
| 4 | ATRX | 224 | KRAS | 366 | SPTA1 | 366 |
| 5 | PIK3CA | 219 | SPTA1 | 362 | KMT2C | 346 |
| 6 | PTEN | 165 | KMT2C | 349 | FAT1 | 338 |
| 7 | CTNNB1 | 154 | ARID1A | 332 | DMD | 326 |
| 8 | PBRM1 | 153 | PTEN | 331 | APOB | 313 |
| 9 | KRAS | 152 | KMT2D | 331 | APC | 290 |
| 10 | ARID1A | 148 | APOB | 318 | ARID1A | 280 |
| 11 | KMT2D | 142 | BRAF | 298 | HUWE1 | 251 |
| 12 | KMT2C | 138 | DMD | 290 | PTEN | 245 |
| 13 | VHL | 131 | FAT1 | 246 | MACF1 | 236 |
| 14 | CIC | 127 | ATM | 236 | NOTCH1 | 222 |
| 15 | SPTA1 | 119 | NF1 | 232 | FBXW7 | 221 |
| 16 | APOB | 115 | MACF1 | 232 | NF1 | 217 |
| 17 | NF1 | 103 | HUWE1 | 227 | RB1 | 211 |
| 18 | SETD2 | 102 | PTPRD | 205 | CREBBP | 209 |
| 19 | ATM | 87 | ZFHX3 | 204 | EP300 | 203 |
| 20 | HUWE1 | 86 | KMT2A | 199 | PTPRD | 201 |
| 21 | DMD | 84 | ERBB4 | 198 | KRAS | 193 |
| 22 | MACF1 | 83 | COL5A1 | 182 | ZFHX3 | 191 |
| 23 | APC | 80 | NCOR1 | 181 | ATM | 190 |
| 24 | ZFHX3 | 79 | MGA | 180 | ATRX | 186 |
| 25 | BAP1 | 76 | KMT2B | 179 | BRCA2 | 184 |
| 26 | FAT1 | 74 | BRCA2 | 175 | NSD1 | 181 |
| 27 | ARID2 | 74 | ATRX | 173 | KMT2A | 180 |
| 28 | CDH1 | 74 | CREBBP | 172 | ERBB4 | 178 |
| 29 | SMARCA4 | 74 | NIPBL | 169 | KMT2B | 174 |
| 30 | PTPRD | 73 | POLQ | 169 | FLNA | 170 |
| 31 | PIK3R1 | 72 | FBXW7 | 165 | POLQ | 168 |
| 32 | NIPBL | 71 | ARID2 | 164 | NIPBL | 167 |
| 33 | NOTCH1 | 71 | CACNA1A | 164 | COL5A1 | 167 |
| 34 | MTOR | 71 | EGFR | 163 | CACNA1A | 159 |
| 35 | CHD4 | 70 | NSD1 | 162 | MED12 | 153 |
| 36 | MGA | 70 | FLNA | 159 | BRAF | 152 |
| 37 | BRCA2 | 66 | CTNNB1 | 157 | CHD4 | 150 |
| 38 | KMT2B | 66 | PIK3R1 | 157 | MGA | 150 |
| 39 | SPOP | 66 | SETD2 | 156 | MYH9 | 149 |
| 40 | NRAS | 66 | SMARCA4 | 155 | SETD2 | 147 |
| 41 | KMT2A | 64 | SETBP1 | 155 | NCOR1 | 145 |
| 42 | COL5A1 | 61 | TAF1 | 154 | CHD8 | 145 |
| 43 | BCOR | 59 | CHD4 | 153 | ATR | 145 |
| 44 | CACNA1A | 59 | MTOR | 153 | USP9X | 143 |
| 45 | USP9X | 58 | MED12 | 152 | EPHA3 | 140 |
| 46 | EGFR | 58 | ARHGAP35 | 151 | ARID2 | 139 |
| 47 | KDM6A | 58 | USP9X | 149 | SMARCA4 | 138 |
| 48 | FUBP1 | 58 | POLE | 147 | KDM6A | 137 |
| 49 | EP300 | 57 | CDH1 | 146 | NFE2L2 | 137 |
| 50 | MED12 | 57 | EPHA3 | 144 | ARHGAP35 | 136 |

Genes are ranked based on the total number of mutations, Total #: Total number of mutations

#### Supplemental Table 3

**Chromosomes with highest number of arm-level gains and losses within each Cell Cycle score subgroup**

| Rank | Cell Cycle Score | | | | | | | | | | | |
|---|---|---|---|---|---|---|---|---|---|---|---|---|
| | Arm-level gains | | | | | | Arm-level losses | | | | | |
| | Low | | Intermediate | | High | | Low | | Intermediate | | High | |
| | Chr | Total # | Chr | Total # | Chr | Total # | Chr | Total # | Chr | Total # | Chr | Total # |
| 1 | 7p | 657 | 20q | 1066 | 8q | 1373 | 1p | 561 | 17p | 1260 | 17p | 1360 |
| 2 | 7q | 606 | 1q | 1056 | 20q | 1113 | 8p | 537 | 8p | 1047 | 8p | 1319 |
| 3 | 1q | 538 | 8q | 1016 | 5p | 1014 | 17p | 523 | 9p | 976 | 4p | 1171 |
| 4 | 8q | 468 | 7p | 980 | 3q | 972 | 6q | 482 | 18q | 965 | 3p | 1136 |
| 5 | 20q | 416 | 20p | 812 | 1q | 914 | 3p | 470 | 16q | 883 | 5q | 1112 |
| 6 | 16p | 415 | 7q | 809 | 7p | 911 | 22 (22q) | 421 | 22 (22q) | 832 | 18q | 1111 |
| 7 | 20p | 359 | 5p | 793 | 20p | 887 | 18q | 408 | 9q | 813 | 9p | 1033 |
| 8 | 5p | 351 | 16p | 488 | 12p | 756 | 9p | 404 | 6q | 784 | 4q | 1020 |
| 9 | 17q | 310 | 3q | 484 | 7q | 629 | 16q | 365 | 18p | 770 | 13 (13q) | 969 |
| 10 | 12p | 308 | 12p | 458 | 18p | 497 | 13 (13q) | 351 | 10q | 747 | 16q | 946 |
| 11 | 12q | 283 | 6p | 442 | 2p | 468 | 9q | 347 | 4q | 728 | 10q | 839 |
| 12 | 16q | 273 | 13 (13q) | 398 | 6p | 455 | 18p | 341 | 13 (13q) | 713 | 11q | 816 |
| 13 | 5q | 268 | 19q | 394 | 19q | 441 | 14 (14q) | 328 | 3p | 696 | 9q | 798 |
| 14 | 17p | 219 | 17q | 375 | 8p | 394 | 21 (21q) | 306 | 4p | 690 | 21 (21q) | 797 |
| 15 | 6p | 208 | 2p | 363 | 17q | 390 | 10q | 291 | 14 (14q) | 669 | 22 (22q) | 793 |
| 16 | 19q | 191 | 10p | 355 | 16p | 372 | 4q | 276 | 5q | 654 | 18p | 764 |
| 17 | 3q | 188 | 8p | 350 | 10p | 371 | 19q | 253 | 15 (15q) | 611 | 11p | 750 |
| 18 | 10p | 170 | 18p | 318 | 21 (21q) | 371 | 3q | 247 | 10p | 596 | 10p | 720 |
| 19 | 19p | 164 | 12q | 312 | 12q | 358 | 11p | 247 | 21 (21q) | 564 | 15 (15q) | 718 |
| 20 | 2p | 163 | 21 (21q) | 273 | 9q | 357 | 4p | 225 | 11p | 555 | 6q | 701 |
| 21 | 8p | 160 | 19p | 251 | 22 (22q) | 327 | 10p | 222 | 1p | 518 | 14 (14q) | 656 |
| 22 | 2q | 155 | 5q | 249 | 9p | 320 | 15 (15q) | 222 | 11q | 471 | 19p | 598 |
| 23 | 21 (21q) | 133 | 2q | 236 | 14 (14q) | 320 | 6p | 217 | 19p | 441 | 1p | 555 |
| 24 | 3p | 129 | 16q | 209 | 13 (13q) | 303 | 11q | 198 | 17q | 318 | 16p | 533 |
| 25 | 18p | 123 | 9p | 208 | 19p | 241 | 2p | 165 | 6p | 304 | 19q | 358 |
| 26 | 13 (13q) | 122 | 14 (14q) | 205 | 16q | 235 | 2q | 162 | 12p | 302 | 12q | 356 |
| 27 | 9q | 112 | 11q | 201 | 11q | 218 | 17q | 158 | 12q | 294 | 17q | 345 |
| 28 | 10q | 104 | 22 (22q) | 196 | 15 (15q) | 217 | 1q | 130 | 16p | 287 | 6p | 335 |
| 29 | 11q | 101 | 11p | 182 | 6q | 215 | 12p | 124 | 19q | 279 | 20p | 321 |
| 30 | 4p | 98 | 9q | 180 | 18q | 209 | 19p | 117 | 20p | 269 | 2q | 315 |
| 31 | 11p | 93 | 4p | 176 | 11p | 206 | 5q | 116 | 3q | 243 | 12p | 297 |
| 32 | 22 (22q) | 90 | 3p | 169 | 2q | 205 | 8q | 94 | 2q | 224 | 5p | 291 |
| 33 | 15 (15q) | 89 | 18q | 159 | 4p | 169 | 12q | 89 | 2p | 209 | 7q | 281 |
| 34 | 18q | 88 | 6q | 155 | 1p | 162 | 20p | 88 | 5p | 188 | 7p | 234 |
| 35 | 14 (14q) | 85 | 15 (15q) | 140 | 17p | 150 | 5p | 83 | 7q | 147 | 2p | 226 |
| 36 | 4q | 74 | 10q | 135 | 3p | 149 | 16p | 58 | 7p | 123 | 3q | 190 |
| 37 | 9p | 72 | 1p | 106 | 5q | 145 | 7q | 42 | 8q | 120 | 1q | 157 |
| 38 | 6q | 65 | 17p | 106 | 10q | 119 | 7p | 23 | 1q | 105 | 20q | 125 |
| 39 | 1p | 28 | 4q | 87 | 4q | 97 | 20q | 23 | 20q | 60 | 8q | 99 |

Total #: Total number of arm-level gains, Chr: Arm-level chromosomal locations, Chromosomal locations were ordered based on the total number of arm-level gains/losses
